# Bound2Learn: A Machine Learning Approach for Classification of DNA-Bound Proteins from Single-Molecule Tracking Experiments

**DOI:** 10.1101/2020.02.20.958512

**Authors:** Nitin Kapadia, Ziad W. El-Hajj, Rodrigo Reyes-Lamothe

## Abstract

DNA-bound proteins are essential elements for the maintenance, regulation, and use of the genome. The time they spend bound to DNA provides useful information on their stability within protein complexes and insight into the understanding of biological processes. Single-particle tracking allows for direct visualization of protein-DNA kinetics, however, identifying whether a molecule is bound to DNA can be non-trivial. Further complications arise when tracking molecules for extended durations in processes with slow kinetics. We developed a machine learning approach, termed Bound2Learn, using output from a widely used tracking software, to robustly classify tracks in order to accurately estimate residence times. We validated our approach *in silico*, and in live-cell data from *Escherichia coli* and *Saccharomyces cerevisiae*. Our method has the potential for broad utility and is applicable to other organisms.

## Introduction

Quantitative information regarding the kinetics of a protein provides valuable insight into the behaviour of the protein, as well as its relationship with other proteins if it is part of a complex. This in turn may inform on the activity of the protein. The residence times of DNA-bound proteins (DBP) can reveal important details on basic cellular processes such as transcription, DNA repair, and DNA replication, at the timescales at which they operate (1-7). This is true for proteins that bind directly at sites on DNA (such as initiator proteins, repair proteins, and chromatin remodellers) and indirectly as part of complexes that bind or translocate on DNA (such as the DNA replication complex (replisome) and RNA polymerase (RNAP)) (1,8).

Recent advances in fluorescence microscopy have allowed us to study protein kinetics directly in living cells, with the most common techniques used being single-particle tracking (SPT), fluorescence recovery after photobleaching (FRAP), and fluorescence correlation spectroscopy (FCS) (3,4,8-13). SPT has the particular advantage of being able to directly observe protein behaviour, allowing for a wealth of information to be extracted from the images, both qualitatively and quantitatively (14,15). Typically, SPT has been used to determine binding kinetics of DBP with very fast kinetics (hundreds of milliseconds to a few seconds), by using capture rates of few to tens of milliseconds. However, many processes operate on much longer timescales posing issues for SPT, including photobleaching and unreliable tracking of single-molecules.

To bypass these issues, a useful approach is to use long-exposure times to blur out diffusing molecules, in combination with stroboscopic illumination to minimize photobleaching (**Figure 1A**). Nonetheless, multiple yet unresolved issues continue to complicate the analysis of this approach. Intensity fluctuations caused by the molecule moving out of focus and photophysics of the fluorophores results in fragmentation of tracks (1,16). Reducing the threshold intensity for spot localization can compensate for this at the cost of introducing false positives. A second common problem is the incorrect assignment of diffusive molecules as DNA-bound, despite motion blurring and tracking parameters to select for only bound molecules. This requires further filtering steps so that only tracks representing true DNA-bound proteins are included in the analysis (1,2,17,18). In contrast to complications associated with tracking algorithms and automated analysis, the user can typically distinguish immobile DBP when looking at the raw images under these imaging conditions as they appear to wiggle around a fix point. This led us to the idea that a machine learning approach would be able to accurately classify the DNA-bound state of a protein.

**Figure 1.**
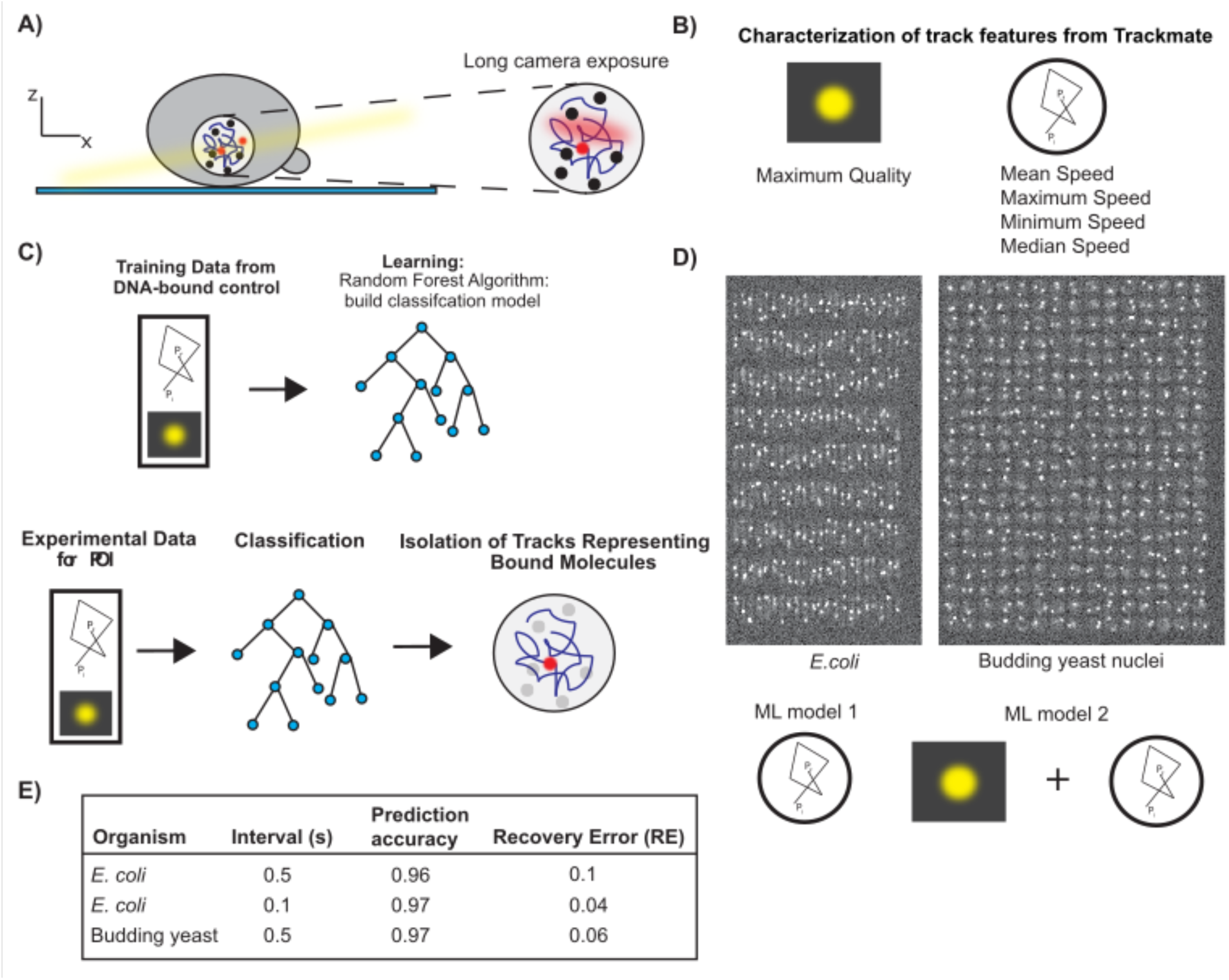
Approach to isolate DNA-bound molecules. A) Diagram of experimental setup to do single-molecule tracking with photoactivatable/photoconvertible fluorophores to calculate residence times on DNA. B) Variables from Trackmate used to predict if a track represents a genuine bound molecule. C) Illustration of the general procedure for ML and how it can be used to predict tracks from experimental data for POI. D) Top - Example images from simulations of single-molecule movies representing *E.coli* cells and budding yeast nuclei. Bottom – Illustration of the variables used in ML models 1 and 2. E) Test data results for different ML models.

Here, we provide a user-friendly and robust method called Bound2Learn to determine residence times of DBP using machine learning and SPT in live *Escherichia coli* (*E.coli*) and *Saccharomyces cerevisiae* (budding yeast), which can be easily extended to other organisms.

## Materials and Methods

### Computer Simulations of Single-Molecule Timelapses

Computer simulations of images were written in Python 3.6.

First, a 5000×5000 array was constructed as an image. Each array element was 10nm, therefore the total size was meant to represent 50µm x 50µm. Cells were placed in a grid-like pattern to prevent overlap and given a low intensity value of around 0.2 per array element to represent cell autofluorescence. *E.coli* cells were modelled as 3D rectangles, with average width of 0.7µm, and an average length of 3µm., while budding yeast nuclei were modelled as spheres of average diameter 2µm. For each cell, unless stated otherwise, two fluorescent molecules were assigned such that their initial locations were confined to the interior of their respective cell. A weighted sampling method was used to assign their initial diffusive state.

To model fluorescent spots, a spot intensity was assigned at the center of the molecules and using a standard deviation of 130nm, the intensity was spread across the region using a Gaussian filter. To represent out-of-focus spots in the budding yeast simulation, the standard deviation used was increased by 5% for every 150nm increase/decrease in z position of the molecule’s position. For example, from +/- 150nm from the z-origin of 0, the molecule was assumed to be in focus. However, if the molecule moved to a |z position| of >150nm but less than 300nm, its standard deviation was increased by 5%. After the Gaussian filter, Poisson shot noise at each element was added by sampling from a Poisson distribution with mean equal to the initial intensity value at that element.

Kinetics for individual molecules were determined using transition matrices: one for the photophysics (e.g. photobleaching), and the other for transitions to different diffusive states. We determined the probability of transition using a time step (τ) of 5ms. We omitted photoblinking as the lower laser powers used in long-exposure experiments typically do not cause it. This simplified the photophysics transition matrix to:

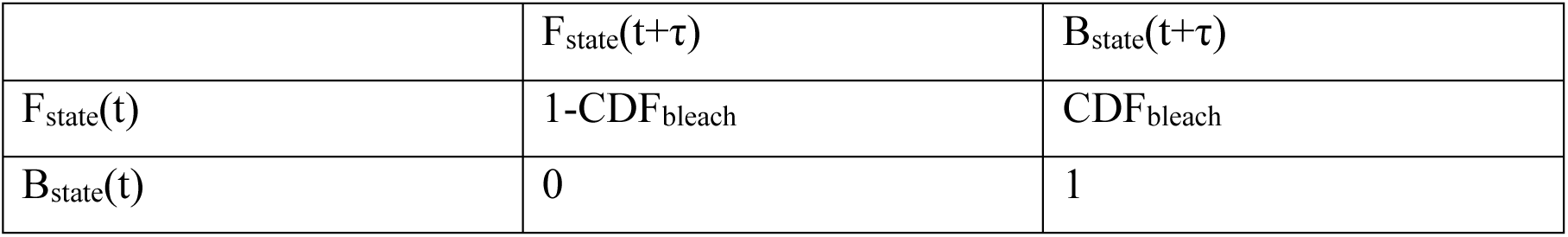

Where the F_state_ is the fluorescent state, B_state_ is the irreversible photobleached state, and CDF_bleach_ is the probability of photobleaching within τ, calculated from the cumulative distribution function (CDF) of an exponential distribution with a specified mean bleach time. If the molecule went to the B_state,_ it was removed from subsequent iterations.

The transition matrix for transitions to different diffusive states was:

| | $Mobile_{state}(t+\tau)$ | $Bound_{state}(t+\tau)$ |
| --- | --- | --- |
| $Mobile_{state}(t)$ | $1 - CDF_{search}$ | $CDF_{search}$ |
| $Bound_{state}(t)$ | $CDF_{bound}$ | $1 - CDF_{bound}$ |

Where Mobile_state_ represents the state with D = D_mobile_, and Bound_state_ represents the state with D = D_bound,_ CDF_search_ is the probability that the molecule switches to the Bound_state_, based on an exponential distribution with a specified mean search time, and CDF_bound_, is the probability that the molecule switches to the Mobile_state_ based on an exponential distribution with a specified mean bound time. In the case of the heterogeneous population of bound molecules, a second Bound_state_ was added with a different bound time.

From the transition matrices and the respective probabilities, we would select the next state using sampling from a multinomial distribution with the weights given by the probabilities.

To simulate molecule movement, the step size in each x,y,z direction was picked from a Gaussian distribution with variance 2Dτ_α_, and α = 1 for the mobile fraction, and 0.4 for the bound fraction to represent subdiffusive behaviour of genomic loci, due to the Rouse model of DNA polymer motion, although we do acknowledge that loci in *E.coli* can undergo more ballistic motions (19-21).

After each time step, the sum of the values of 10×10 array elements was taken to simulate a 100nm pixel size and get a 500×500 pixel image. Subsequently, camera noise was added by sampling from gaussian distribution with mean = 150 and standard deviation = 20. To obtain images of different exposure times, time steps were integrated (e.g. 100ms = 5ms * 20 steps of integration). In the case of time intervals, we allowed molecule movement but no image formation until the next image is taken. E.g. for 1s interval, a 500ms image was followed by 500ms of only molecule movement but no image formation, followed by another 500ms image.

### Construction of Top2 and TBP HaloTag fusions

Strains used in this study are all from a BY4741 background and are shown in **Table S5**. Plasmids used in this study are derivatives of pUC18 (ColE1 origin, Ampicillin resistance); pTB16 carries mNeonGreen and a downstream NatMX marker, while pSJW01 carries the HaloTag gene with a HygB marker. Both mNeonGreen and HaloTag genes encode an 8 amino acid linker at the 5’ end (sequence: GGTGACGGTGCTGGTTTAATTAAC). Plasmids were maintained in *E. coli* DH5α and were extracted by growing in LB with 100 μg/ml ampicillin then using the Presto Mini Plasmid Kit (Geneaid).

All PCRs were made using either Phusion or Q5 (NEB). PCR reaction were in a volume of 50 μl and included water, 3% DMSO, the reaction buffer, 2.5 mM of each dNTP, 0.2 μM of each primer, either 1ng of plasmid DNA (for insertions) or 1 μl of genomic DNA (for screening insertions), and 0.5 μl of polymerase.

Fluorescent fusions were made by PCR amplification from pTB16 or pSJW01 using the primers listed in **Table S6**. PCR products were transformed into wild-type diploid BY4743. A single colony was grown at 30°C in 5 ml yeast peptone dextrose (YPD) overnight, then diluted to a final OD_600_ of 0.1 in 10ml of YPD. Cells were taken at OD_600_=0.5-0.6 and centrifuged at 4000 rpm for 5 min. The pellet was washed twice with 25 ml of sterile deionized water, then once with 1ml of 100mM lithium acetate. Cells were then resuspended in 240 μl of 50% PEG, then 50 μl of salmon sperm DNA (thawed at 95°C for 5min, then incubated on ice for at least 10min), 50 μl of the PCR product and 36 μl of 1M lithium acetate were added in this order. The mixture was thoroughly mixed by pipeting and incubated on a rotator at 25°C for 45min, followed by heat shock at 42°C for 30min. Cells were pelleted in a microcentrifuge, washed in 500 μl of sterile water, then resuspended in 200 μl of YPD and plated on YPD agar. After growing at 30°C overnight, the cell lawn was replica-plated onto selective YPD agar, either with 100 μg/ml cloNAT (Werner) for mNeonGreen, or with 200 μg/ml Hygromycin B (Life Technologies) for HaloTag. Transformants were screened for the presence of an insert by PCR using the indicated primers in **Table S6**.

Confirmed clones were then sporulated by taking 750 μl of a YPD overnight cultures, washing 4 times with 1 ml sterile deionized water, then washing once with 1 ml of potassium acetate sporulation medium (Kac), and finally resuspending in 2 ml of KAc and incubating at 25°C wish shaking. After 5 days the sporulating cultures were checked by microscopy for the appearance of numerous tetrads, then 750 μl was taken and washed 3 times in sterile water before final resuspension in 1ml water and storage at 4°C. For dissection, 45 μl of spores were treated with 5 μl of zymolase for 10 min, then tetrads were dissected on YPD plates to isolate haploids with the tagged fusion. Genomic DNA was isolated from the haploid by vortexing the cells in the presence of zirconia/silica beads, followed by phenol extraction and ethanol precipitation. The insertion site was once again amplified using the same screening primers as above, and the PCR product was sequenced to confirm that the tag and linker were both mutation-free.

The Top2-Halo and Spt15-Halo (TBP, TATA-Binding Protein) haploids were combined with PCNA-mNeonGreen (from YTB31) and the pdr5Δ::KanMX deletion (from a haploid sporulated from YTK1414) by mating. To do this, 10 μl of water was spotted on a YPD plate, and colonies from the Mat a and Mat α haploids to be combined were mixed together into the water drop and incubated at 30°C overnight. Cells were then restreaked on an auxotrophic -Met-Lys plate on which only the mated diploid could grow. Diploids resulting from a mating were dissected as above. PCNA-mNeonGreen (from YTB31) and the *pdr5Δ::KanMX* deletion (from a haploid sporulated from YTK1414) were combined by mating and the resulting diploid was dissected to isolate strain ZEY098, a haploid with both PCNA-mNeonGreen and *pdr5Δ::KanMX*. ZEY098 was then mated with either a Top2-Halo or Spt-15-Halo haploid of the opposite mating type, and the resulting diploids were dissected to create haploids with all three markers (HaloTag, PCNA- mNeonGreen and *pdr5Δ*) that were then used for imaging and control experiments. The genotypes of these imaging haploids are detailed in **Table S5**.

### Single Molecule Imaging in Budding Yeast

A single colony from a YPD plate was placed in 5mL synthetic complete (SC) medium and grown with shaking at 30°C for ∼5-6 hours. This culture was diluted by transferring ∼50uL into 5mL of fresh S.C and grown overnight at 30°C. The overnight culture was diluted to 0.15 the next day and grown until the optical density (OD) reached 0.30. 1mL of this culture was spun down for 1 min @ 4000RPM, and the pellet was resuspended in 500uL of fresh S.C. Janelia Farms photoactivatable 549 (JF-PA549) was added to the 500uL culture for a final dye concentration of 50nM, except for YTK1434-Halo (Histone H3), where a concentration of 10nM was used to compensate for the higher copy number. This culture was placed in a thermomixer at 30°C and 500RPM for 40 minutes. After incubation, 3 wash cycles using fresh S.C were done to wash away unbound dye. After the final wash step, the pellet was resuspended in 50uL of S.C, and 3uL of the culture was placed on an agarose pad consisting of SC and Optiprep (Sigma), within a Gene Frame (Thermo Scientific). The pad was made by taking a 2% agarose Optiprep mixture (0.02g in 1mL Optiprep) - that was heated to 90 degrees - and mixing 500uL with 500uL 2xSC, resulting in a 1% agarose 30% Optiprep SC mixture. Approximately 140uL of this mixture was placed within the Gene Frame, with excess being removed with a KimWipe. Prior to imaging, we waited ∼15 minutes to let any unbound dye be released.

Coverslips were cleaned with the following steps: 1) Place in 2% VersaClean detergent solution overnight. 2) Wash with MilliQ water 3x. 3) Sonicate in acetone for 30 minutes. 4) Wash with MilliQ water 3x. 5) Place in methanol and flame coverslips using Bunsen burner. 6) Place in Plasma Etch plasma oven for 10 minutes.

Microscopy was done at 23°C, on a Leica DMi8 inverted microscope with a Roper Scientific iLasV2 (capable of ring total internal reflection fluorescence (TIRF)), and an Andor iXON Ultra EMCCD camera. An Andor ILE combiner was used, and the maximum power from the optical fiber was 100mW for the 405nm wavelength, and 150mW for the 488nm and 561nm wavelengths. The iLasV2 was configured to do ring highly inclined and laminated optical sheet (HILO), for selective illumination and single-molecule sensitivity. Metamorph was used to control acquisition. A Leica HCX PL APO 100x/1.47 oil immersion objective was used, with 100nm pixel size. Any z-stacks were doing using a PInano piezo Z controller.

Single-particle photoactivated localization microscopy (sptPALM) experiments were performed by activating molecules with low power (0.5% in software) 405nm light to photoactivate ∼1 molecule/cell, followed by stroboscopic, long-exposure (500ms) illumination with 561nm light (5% in software) to image primarily bound molecules. A brightfield image and a z-stack of 6um (0.3um step size) in the 488nm channel, was taken before and after each timelapse, to ensure normal cell health and to find nuclei.

### Tracking Analysis

Tracking was done with Trackmate (22). Spots were localized using the Laplacian of Gaussian (LoG) method, with an estimated spot size of 2.5 pixels, with the exception of the *E.coli* experimental data where it was set to 5 pixels. The intensity threshold was set a bit lower to prevent track fragmentation due to intensity fluctuations. The linear assignment problem (LAP) algorithm was used to form tracks with costs on quality ranging from 0.1-0.5. We set a gap frame of 1 to allow temporary disappearance of the molecule, and track merging and splitting was allowed in cases where multiple molecules crossed paths with one another.

To isolate tracks found only in cells/nuclei, we used the binary images to locate tracks whose mean positions coincided with values of 1 in the binary image.

### Machine Learning and Tracking Analysis

All machine learning and subsequent analysis for estimation of residence times was done using Matlab.

To construct training data sets, we had binary classification, with a value of 0 assigned to false positive/diffusing molecule, and a value of 1 to a track representing a bound molecule. We manually looked at the raw image data to determine if the molecule appeared immobile.

For the learning procedure, the “TreeBagger” function in Matlab was used, representing the random forest algorithm. The hyperparameters that were adjusted were: InBagFraction (representing the fraction of the training data given to each tree), MinLeafSize (minimum leaf size), NumPredictorstoSample (number of predictors to sample at each node), and NumTrees (the number of trees to construct). The hyperparameters were adjusted until the best OOB error was achieved.

For GMM fitting, the expectation-maximization (EM) algorithm was used.

After the final classification, we analyzed the tracks to extract residence times.

We fit the track durations of the resulting tracks with a truncated exponential model, to compensate for discarding short duration tracks, using Maximum Likelihood Estimation (MLE) through Matlab’s “mle” function, to calculate the mean track duration.

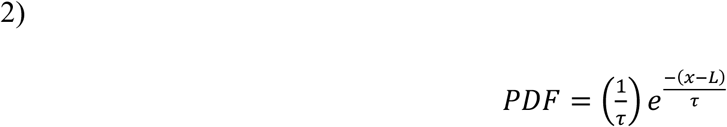

where τ is the mean track duration, and *L* is the truncation point. For photobleaching controls, this was equivalent to estimating the mean bleach time.

The 95% confidence intervals were calculated by bootstrapping 1000 samples.

In cases where the experimental data was taken with a longer time interval than the photobleaching control, we used the following equation to calculate *Tbleach h*(1,23):

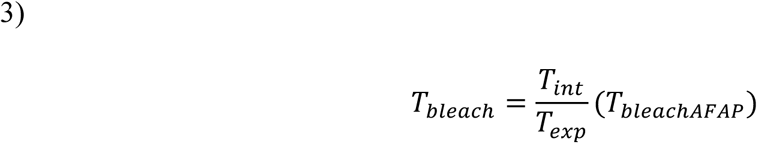

Where *T*_*int*_ and *T*_*exp*_ are the time interval and exposure, respectively, used to acquire the data, while *T*_*bleachAFAP*_ is the photobleaching time for data collected with continuous exposure, which in in the case of the data used here, would represent 500ms interval

Bound times were calculated using the following equation, after combining data from multiple experiments collected with the same time interval:

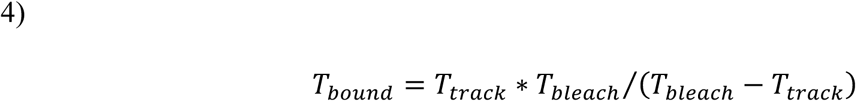

To calculate the errors on the estimate, we performed bootstrap sampling on the track durations to the following equation:

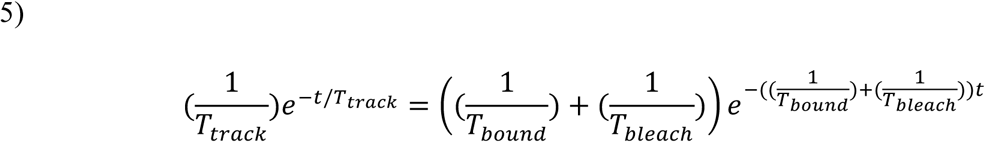

With 10% variation allowed for the *T*_*bleach*_ estimate, in order to obtain biologically sensible results.

To check for two-exponential mixtures, the track durations were fit with the following two-exponential model:

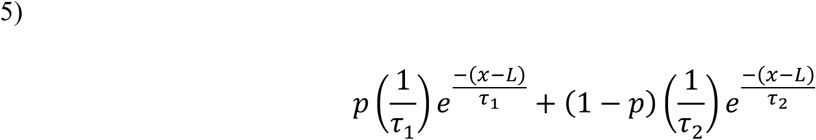

where τ_*1*_ = (*Tbleach + Tbound*_*γ*_)/(*Tbleach* \**Tbound*_*γ*_), τ _*2*=_ = (*Tbleach + Tbound*_*Ψ*_)/(*Tbleach * Tbound*_*Ψ*_), is the mixture proportion, and *L* is the truncation point.

The lower and upper bounds on the two binding timescales were 0.0001s and 6000s, respectively, while allowing for a 10% variation in the bleaching estimate.

To check for overfitting and to identify whether the two-exponential model significantly fit the data better, we used the BIC test and the Loglikelihood ratio (LLR) test, as described in (1). We looked for cases when the two-exponential model estimates did not simply return the lower and/or upper bounds as this would indicate that no physically sensible solution was found.

### Diffusion Coefficient Estimation

Diffusion coefficients were estimated from individual tracks by calculated the slope of the time lag (τ) vs mean squared displacement (MSD) curve. The equation used was modified from(24):

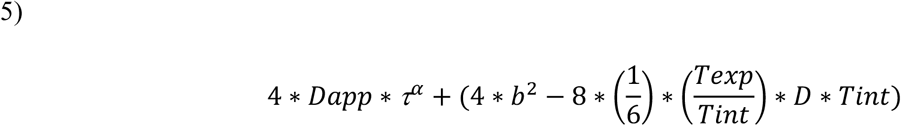

Where *Dapp* is the apparent diffusion coefficient, α is the anomalous diffusion constant, and *b* is the static localization error. We used this equation to correct for dynamic localization error due to the molecule moving within the exposure time, which in our case was quite long.

To classify tracks using their Dapp estimates, we performed GMM fitting on the distribution of Dapp, with 2 components, and clustered tracks according to their Dapp to assign them as either being in the bound state vs diffusing/noise. Clustering was done by first calculating posterior probabilities to each track (the probability it belongs to a diffusive state given its Dapp), and then selecting the diffusive state with the maximum posterior probability.

To gain a better estimate of the diffusion coefficient for Top2, TBP, and Histone H3, we performed a weighted, least -squares fit using the MSD values averaged over the tracks, after they were classified by Bound2Learn. We did not include tracks with a goodness of fit (gof) score of less than 0.70 from the previous step, given the noisiness of some of the MSD curves. We also included lower and upper bounds on the estimates to ensure physically sensible estimates, and in cases where on the estimates was the lower or upper bound value, we discarded data points at higher τ values. This is a known issue given there are fewer data points in the calculation of MSD at the higher τ values leading to more noisier traces.

## Results

### Random Forest for single-molecule tracking classification

Machine learning (ML), including its branch of deep-learning, is a powerful tool for image analysis and classification, with the most common implementation being supervised learning, whereby a labelled training data set is given to the ML algorithm, which then builds a model for subsequent classification (25,26). Our motivation arose from recognizing the limitations associated with automated detection and tracking, and our desire to develop a classification method to compensate for these limitations. We recognized that a known DBP (e.g. histone H3) could be used to construct a training data set manually, for how DBP in general should move and how single-molecules should look like in images, from which we can build a ML model. We can also use this control as a photobleaching control ? if it exhibits stable binding ? when we estimate the residence time for our DNA-bound protein of interest (POI) (1).

We used Trackmate, a freely available plugin in Fiji, to track molecules and used variables from the tracking output that we believed to be good predictors of single-molecule DBP and reduced the cross-validation error (CVE) (**Figure 1B**) (22). To predict whether a molecule moves like a DBP we used the following variables from the tracking output: mean speed, maximum speed, minimum speed, and median speed. For predicting whether the track represented a genuine molecule, we used the maximum quality variable – a parameter corresponding to the intensity of the molecule as well as its shape (22). Classification of the track was done manually by looking at the raw image data, with binary classification: 0 for a diffusing molecule or noise, and 1 for a track representing a genuine immobile, DNA-bound molecule. We chose to use the random forest algorithm to construct a model from the training data, as it is accurate, tolerant of noise, and less prone to overfitting (27) (**Figure 1C**). The various parameters used to construct the different random forest models are listed in **Table S1.** We used this model to classify tracks for our POI (Figure 1C), and subsequently determine their residence time after correcting for photobleaching.

We first tested our approach using computer simulations of single-molecule timelapses of *E.coli* and nuclei of budding yeast (see **Materials and Methods**). The main difference between the two simulations aside from cell shape, was that the budding yeast simulation had a distortion in the shape of the molecule based on its position in z, to model the point spread function (PSF) and emulate the molecule going out of the focal depth of the objective. Under our experimental conditions, *E.coli* cells have diameters of 0.7μm, which is not far from the estimated focal depth of high numerical aperture objectives commonly used in single-molecule studies (∼0.4 μm), so we assumed molecules to be in focus, regardless of their z position(1). We first constructed training data sets from simulated data with 500ms exposure, no time interval, and a mean bleach time of 10s (**Table S2)**. We had a stable, bound fraction (Diffusion coefficient (D) of D_bound_= 0.005μm^2^/s, bound time >>> bleach time), along with a mobile fraction (D_mobile_ = 0.5μm^2^/s) **(Figure 1D).**

We only considered tracks with ≥ 4 localizations, as it is difficult to discern the state of the molecule for shorter tracks. In order to make use of a single training data set, instead of constructing multiple training data sets for classifying data sets collected under different conditions, we constructed two models: ML model 1 only has the speed variables and ML model 2 has the speed variables along with maximum quality (**Figure 1D)**. We calculate the mean of ‘mean speed’ as well as the mean of ‘maximum quality’ from the tracks classified as being bound in the training data and used these values to scale the speed and quality variables, respectively, for data collected with different time intervals and/or illumination intensities **(Figure S1).**

The entire procedure for the classification of tracks for a POI is as follows:

1. We performed a two-component Gaussian mixture model (GMM) fit on the log of mean speed, of all the tracks, and the component with the lowest mean is selected as representing the immobile molecules. This step does not need to be robust as it is only used to do some initial filtering for subsequent steps.
2. Using this value, we scale the speed variables accordingly, using the mean of ‘mean speed’ calculated from the training data, as follows: scale factor_speed_ = mean (mean speed_training_)/mean(mean speed_boundPOI_). This helps to ensure the ML models can classify tracks obtained from different time intervals than the training data. We then run the tracks through ML model 1 for initial filtering.
3. From the resulting tracks, we calculate the mean of ‘maximum quality’ using a two-component GMM, similar to step 1 except selecting the component with the higher mean and use the same variable from the training data to scale the quality variable, similar to the previous step. We then run the tracks through ML model 2 for final classification.

To assess how well the procedure worked, we quantified both the accuracy (proportion of tracks accurately predicted to be bound), and the recovery error (RE) (the fraction of tracks known to be representing bound molecules that were missed by the classification procedure). While high classification accuracy is important to ensure accurate estimation of residence time, we also wanted to make sure that the recovery error is small, as single-molecule studies are often plagued by low sample sizes thereby reducing confidence in the estimate (1,14). In addition, the out-of-bag (OOB) errors, equivalent to the CVE for random forests (27), were estimated for each ML model **(Table S1)**. On test data, we obtained an accuracy of 0.96 and RE of 0.10 for the *E.coli* simulation, and an accuracy of 0.97 and RE of 0.06, for the budding yeast simulation (**Figure 1E)**.

### Accurate estimation *in silico* of residence times under different conditions

Next, we tested how well the ML models would work on simulations of image data with different time intervals and spot intensities (**Materials and Methods, Table S2)**. First, we tested on data with a 1s time interval and the same spot intensities as the training data set (**Videos S1 and S2)**. The mean residence time was set to 8s, while the bound fraction was set to 0.5. As **Figure 2A** (top) illustrates, we first calculate the mean of ‘mean speed’ of the bound population, in order to scale the speed variables. After the final classification step, for both the *E.coli* and budding yeast simulations, we were able to obtain a mean residence time estimate (∼7s) that was in close agreement with the 8s set in the simulation (**Figure 2A** (bottom)). In addition, the accuracy values were 0.93 and 0.96, with recovery errors of 0.11 and 0, respectively, for the *Ecoli* and budding yeast simulations (**Table S3)**.

**Figure 2.**
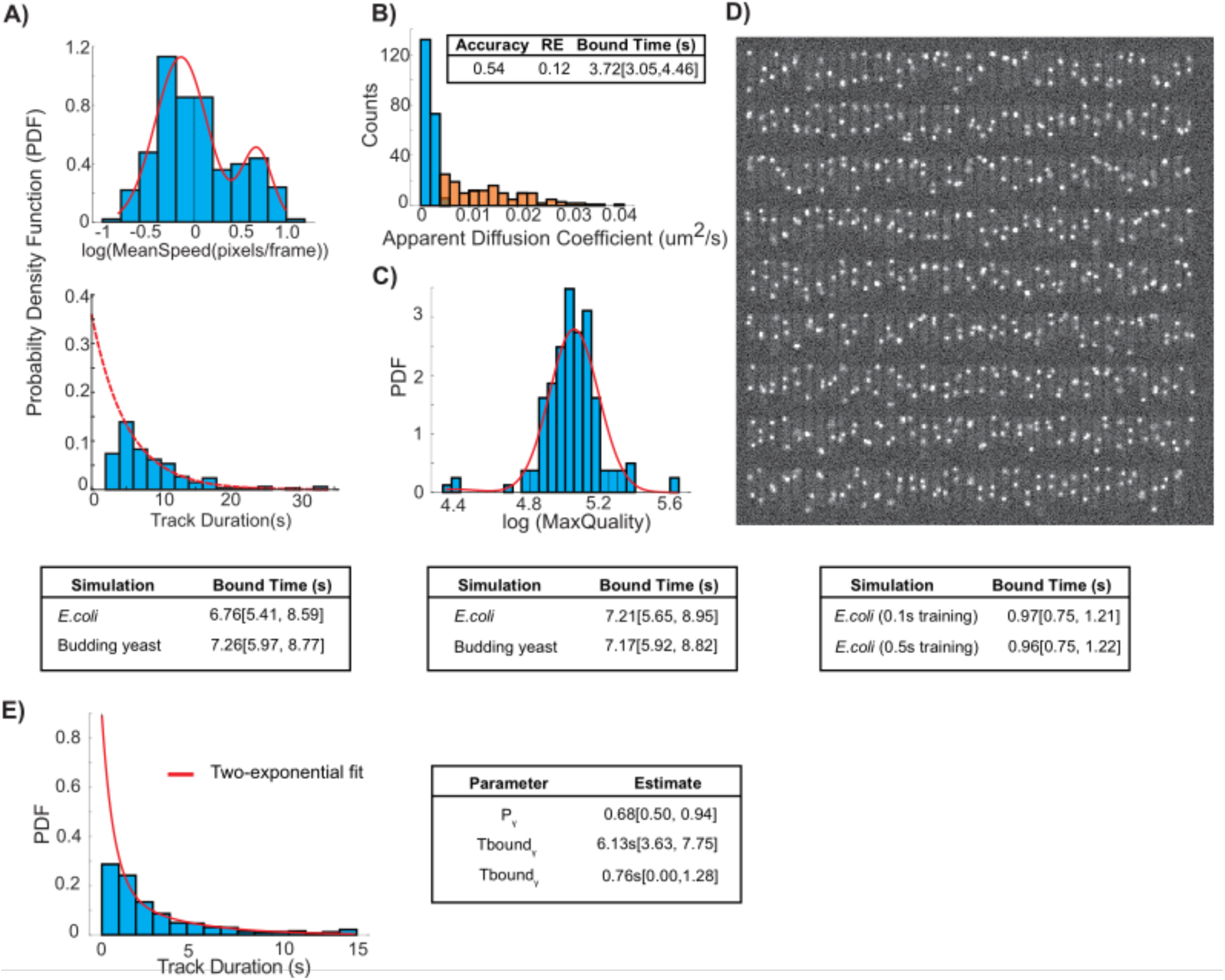
Estimating residence times over a range of conditions. A) Top – Diagram illustrating the GMM fit to determine the mean of mean speed for 1s interval data, which can be used to rescale the speed variables. Bottom – Estimates of residence times in both simulations (N = 169 tracks for *E.coli* and N = 232 tracks for budding yeast). B) Top – Diagram illustrating the GMM fit to maximum quality values from a data set with lower fluorescent intensities. Bottom – Estimates of residence times from simulated data representing poorer image quality (N = 159 tracks for *E.coli* and N = 227 tracks for budding yeast). C) Distribution of apparent diffusion coefficients from *E.coli* simulation, with GMM fitting, followed by clustering to isolate tracks representing bound molecules (blue) and diffusing/noise (orange). N = 291 tracks. D) Top – Example of image from simulated 100ms timelapse in *E.coli*. E) Two-exponential fitting to a data set with heterogeneous population of bound molecules (N = 508 tracks). For all estimates, 95% confidence intervals are shown.

With the *E.coli* simulation, we also tested the accuracy and RE by using a more traditional approach of estimating apparent diffusion coefficients (Dapp) through mean-squared displacement (MSD) analysis, GMM fitting to determine diffusive fractions, and assigning individual tracks to a diffusive state using a cluster method (**Materials and Methods**). We observed that there was a large spread in the apparent diffusion coefficients, consistent with the difficulty of obtaining precise estimates from single tracks using MSD analysis, given that they are inherently noisy (24) (**Figure 2B**). Furthermore, the accuracy and RE using this approach was 0.54 and 0.12, respectively, with a bound time estimate much lower than 8s, suggesting a significant fraction of tracks resembling diffusing molecules or noise were being classified as bound (**Figure 2B**). This suggests that while it is possible that mean values of the Dapp for different states may be accurate, due to precision issues from MSD analysis, assigning individual tracks to different states based solely on their Dapp can be error prone. We then tested whether we could obtain an accurate estimate in a situation representing poorer image quality, by lowering the integrated spot intensity from 3000 to 2000. As **Figure 2C** (top) illustrates, we use the GMM fit on the max quality values to find an appropriate scaling factor. After classification, we find that we can still recover an estimate of the residence that is within error of the known value, for both simulations, with accuracy values and REs similar to the previous condition (**Figure 2C, Table S3)**. In contrast, the MSD analysis approach on the same data resulted in a lower accuracy of 0.79, with a RE of 0.096, and bound time estimate of ∼5 s (**Table S3**).

While a long exposure of 500ms helps to blur out diffusing molecules, it also restricts the temporal resolution achievable, and makes it difficult to obtain precise estimates for faster processes, as fewer tracks will remain if using a track localization acceptance threshold. Therefore, we constructed a training data set with 100ms exposure (accuracy and RE on test data was 0.97 and 0.06, respectively (**Figure 1E**)), and tested on data with a mean residence time set to 1s while the bleach time for this condition was set to 2s (20 frames) (**Figure 2D, Video S3)**. Once again, we were able to obtain an estimate (∼0.97s) that was in agreement with the 1s residence time, although with lower accuracy and higher recovery errors than with 500ms, as one would expect under conditions where the separation between two diffusive states is more difficult to discern. (**Figure 2D)**. We tested this by changing D_mobile_ to 5 μm^2^/s, and found the errors were drastically reduced, with an accuracy of 0.98 and RE of 0.08 **(Table S3)**. We then asked whether the ML models built with the 500ms training data can be used to classify the 100ms data, and surprisingly, we found that it still performed well (**Figure 2D)**, with comparable errors to the 100ms ML models (accuracy = 0.86 and RE = 0.16 with 500ms ML model, vs accuracy = 0.85 and RE = 0.18 with 100ms ML model) (**Table S3)**.

We also found that we could get accurate estimates of the residence times of a heterogeneous population of bound molecules, where two distinct binding regimes were present: mean Tbound_γ_ set to 7s while mean Tbound_*Ψ*_ set to 1s (**Figure 2E).** We note that for the simulation representing a heterogeneous population of bound molecules, we had to change the bleach time to 10s, in order to recover the two binding times. As others have alluded too, detecting multiple populations is highly dependent on acquisition settings (28). These results show that one can use ML models constructed from a single training data set and use them for accurately classifying tracks obtained from data collected under widely different conditions, along with obtaining accurate residence times of homogeneous and heterogeneous bound populations. It is important to highlight that this analysis does not require additional steps to identify appropriate thresholds to filter out noise and/or diffusing molecules, which can be time-consuming and a source of heterogeneity in the analysis.

### Experimental validation in *E.coli*

We then determined if Bound2Learn could estimate residence times from imaging of *E.coli*. Unless stated otherwise, estimates were obtained by fitting track durations using a truncated exponential model (we discarded tracks with <4 localizations as they were not reliable) to data from multiple experiments collected under the same acquisition settings to obtain higher statistical confidence (**Materials and Methods, Table S4**). We reanalyzed a subset of the data reported in (1) to test if we could obtain consistent results. For our DNA-bound control we used LacI, a transcriptional repressor, fused to the photoconvertible mMaple, which upon illumination with 405nm light, converts to a red fluorescent form (29). LacI has been reported to bind stably to the *lacO* array site on DNA (∼5 minutes), thus making it suitable as a photobleaching control, under our acquisition times (30) **(Figure S2)**. Although we had initially used this strain for photobleaching correction in (1), we decided to use it for constructing a training data set as well. The LacI data used for the training data was collected with 500ms exposure and continuous acquisition, resulting an estimated photobleaching time of 13.60s **(Figure 3D)**.

**Figure 3.**
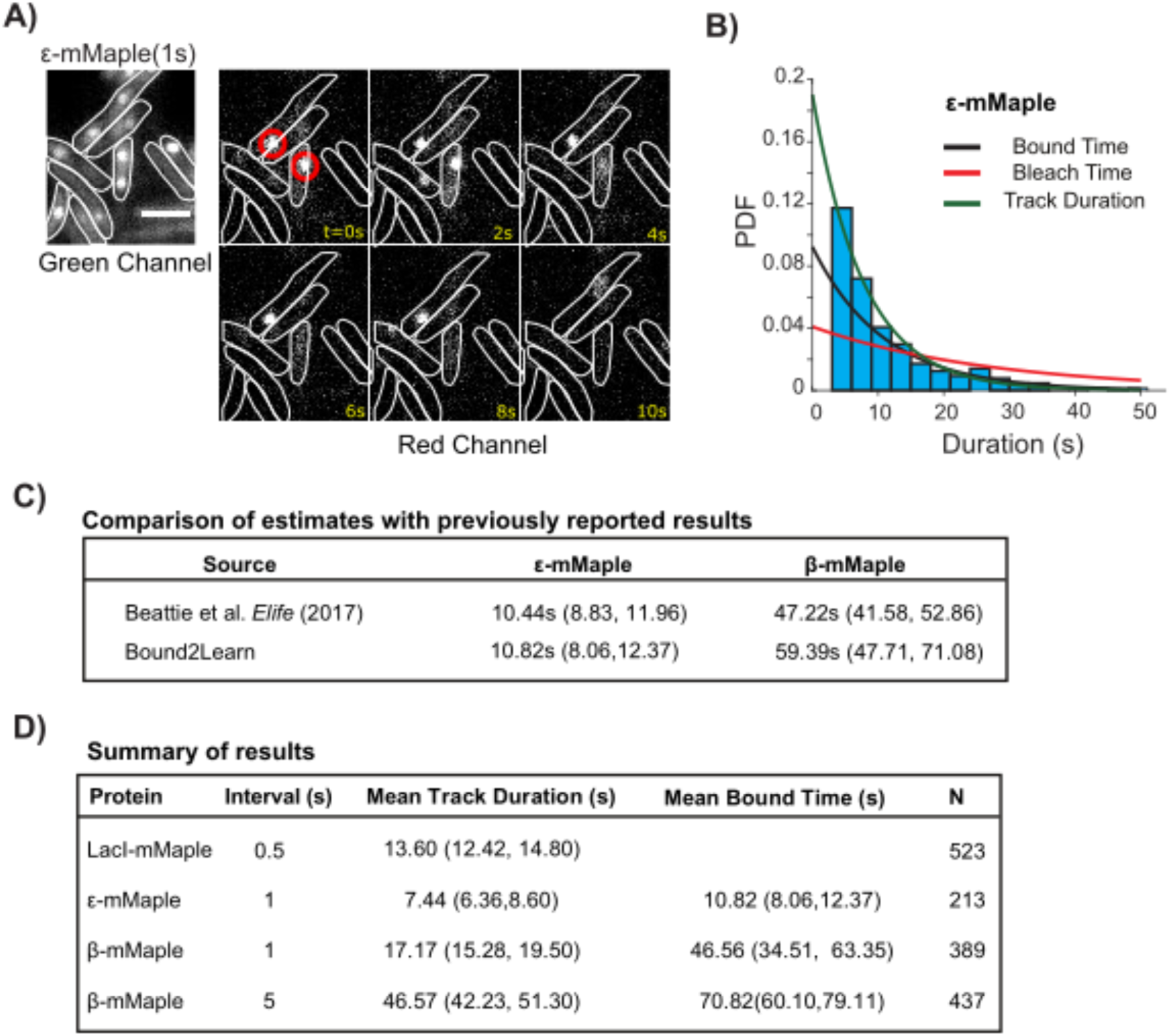
Estimating residence times for PolIII subunit, ɛ. A) Example of dnaQ-mMaple timelapse. Red circles indicates molecules that were classified as being bound with Bound2Learn. Scale bar = 2um. B) Histogram and fit of track durations from combined data set of ε. C) Estimation of residence times and comparison to estimates from (1). Estimate for β was a weighted average calculated from bound times estimated with different time intervals. D) Summary of results showing both mean track duration estimates and mean bound time estimates from combined data sets. 95% confidence intervals are shown next to estimates.

Once the ML models were constructed, we asked whether we could get an accurate estimate of the residence time for *E.coli* DNA polymerase, PolIII, for which we had previously reported to have a residence time of around ∼10s, (1). We reanalyzed a dataset from (1) taken of the PolIII subunit, ε, also tagged with mMaple (dnaQ-mMaple), to determine if Bound2Learn could give similar results (**Figure 3A)**. Our analysis resulted in an estimate of 10.82s (8.06, 12.37s), in very close agreement with the previously reported estimate **(Figure 3B and 3C)**. We also reanalyzed data taken of the DNA binding clamp, β_2_ (dnaN-mMaple), which we previously reported to have a bound time of 47.22s (1). As mentioned before, stable proteins pose an issue for tracking as intensity fluctuations (e.g molecule moving out-of-focus) can result in track fragmentation (1). Bound2Learn is less sensitive to such intensity fluctuations since it allows the use of lower intensity thresholds during the localization step and still result in accurate classification. We analyzed data collected with 1s and 5s time intervals, and then calculated a weighted average as our final estimate for mean bound time. Our estimate using Bound2Learn was 59.39s (47.71s, 71.08s) is slightly higher, but still within error, than our previously published estimate (**Figure 3C**). Use of a different localization and tracking software than the one in our previous publication may also explain part of this difference (1). Our full results are summarized in **Figure 3D**, showing the estimates for mean track durations and mean bound times for the different proteins, and different time intervals in the case of β_2._ Overall, these results suggest that Bound2Learn robustly estimates residence times of proteins with different binding behaviours in *E.coli*.

### Estimating Residence Time of Topoisomerase II in Budding Yeast

We then asked if our approach could be used in live haploid budding yeast. For our photobleaching control and training data construction, we used histone H3 fused to HaloTag (H3-HaloTag), due to its expected long residence time and high bound fraction (31) (**Figure S2 Table S5)**. In order to detect the protein, we incubated cells with the cell-permeable photoactivatable (PA) dye, JF- PA549 (32). The experimental protocol was very similar to that described in *E.coli* (1), consisting of the use of highly-inclined and laminated optical (HILO) sheet stroboscopic illumination with 500ms exposure, but with cycles of low-dose 405nm activation every 40 frames **(Materials and Methods)**. To improve image quality with budding yeast, we also used the refractive-index matching media, Optiprep, to minimize light refraction due to the cell wall (33). The mean photobleaching duration estimated, with 1s interval acquisition, was ∼22 seconds **(Table S4, Figure S2)**.

As experimental test, we tagged Topoisomerase II (Top2) with HaloTag (Top2-HaloTag) and to segment nuclei in order to isolate tracks found only in S-phase nuclei, we tagged proliferating nuclear cell antigen (PCNA) with mNeonGreen (Pol30-mNeonGreen) and acquired a z-stack in the green channel prior to acquisition (**Table S5)**. The resulting z-stack went through a smooth manifold extraction (SME) process for better image quality of the nuclei (34). We chose Top2 as mammalian Top2 enzymes have been reported to be dynamic in interphase cells, with FRAP t_1/2_ lifetimes of 1.8-10s, although no precise residence time has been determined (13). We collected the data under the same acquisition settings as H3-HaloTag. As observed in **Figure 4A**, we were able to visualize dissociation events of the protein under our acquisition settings. Following tracking analysis and ML classification, we obtained an estimate for the residence time of Top2 of ∼30s, consistent with dynamic behaviour reported previously_[RRD1]_ in mammalian cells (**Figure 4B and 4C, Table S4)**. We also performed similar experiments in a strain lacking a HaloTagged protein and could not observe any molecules from the timelapses, and in addition, no bound tracks were obtained after classification with Bound2Learn (**Figure S3**). Therefore, nonspecific binding of the dye is unlikely contributing to our estimates. It should be noticed that despite slightly poorer spot quality compared to *E.coli* (compare **Figure 3B vs Figure 4B)**, Bound2Learn was able to obtain estimates in this system, suggesting our approach can be used across a range of imaging conditions.

**Figure 4.**
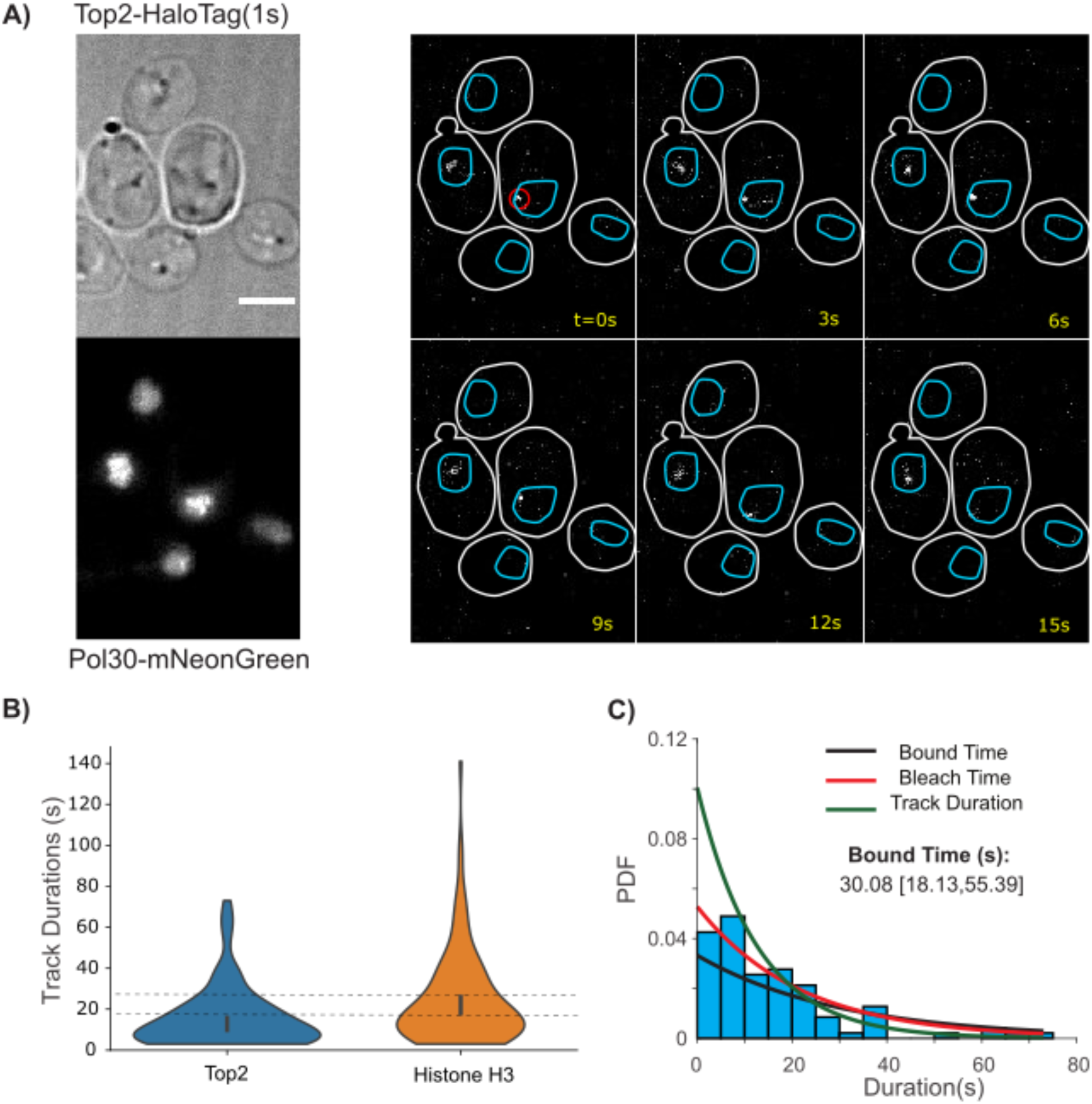
Estimating residence time of Top2. A) Example of Top2-HaloTag timelapse after photoactivation. Red circle indicates molecule classified as being bound with Bound2Learn. Also shown is the SME of a Pol30-mNeonGreen z-stack. Scale bar = 3um. B) Violin plots of track durations for Top2 (N = 94), and Histone H3 (N = 123). Error bars represent 95% confidence intervals. Dotted horizontal lines represent upper and lower bounds of the 95% confidence interval of Histone H3. C) Histogram and fit on Top2 track durations. Note that bin counts for short durations are smaller than expect given that short tracks of <4 localizations were discarded. This was compensated for during the fitting procedure (Materials and Methods). Errors are represented by 95% confidence intervals

To further confirm that our approach works with single-molecule data from budding yeast, we performed similar experiments with a HaloTag fusion of TATA binding protein (TBP) (SPT15- HaloTag), a general transcription factor, which has been shown in budding yeast to have FRAP recovery times < 15s, and with a minimal immobile fraction, suggesting it is dynamic on DNA, although the exact binding time has not been determined (8) **(Table S5).** We collected the data with continuous exposure (500ms intervals) rather than with a 1s time interval, to prevent the possibility of TBP binding to neighboring binding sites thereby artificially increasing its estimated residence time. Our data confirms the dynamic behaviour of TBP, as we observed dissociation within a few seconds (**Figure 5A, Video S4**). Few of the spots detected were in S-phase nuclei, indicating that some transcription (e.g. expression of histones) occurs during S-phase (35). Our measured photobleaching time using histone H3 was 12.19s (**Table S4**). When we quantified the track durations for TBP, we observed a significantly different track duration distribution relative to histone H3, with an abundance of short duration tracks, suggesting most tracks ended due to dissociation of the protein as opposed to photobleaching (**Figure 5B, Table S4**). Our estimate for mean bound time for TBP, after photobleaching correction, was 11.43s, consistent with TBP being dynamic on DNA (**Figure 5C**).

**Figure 5.**
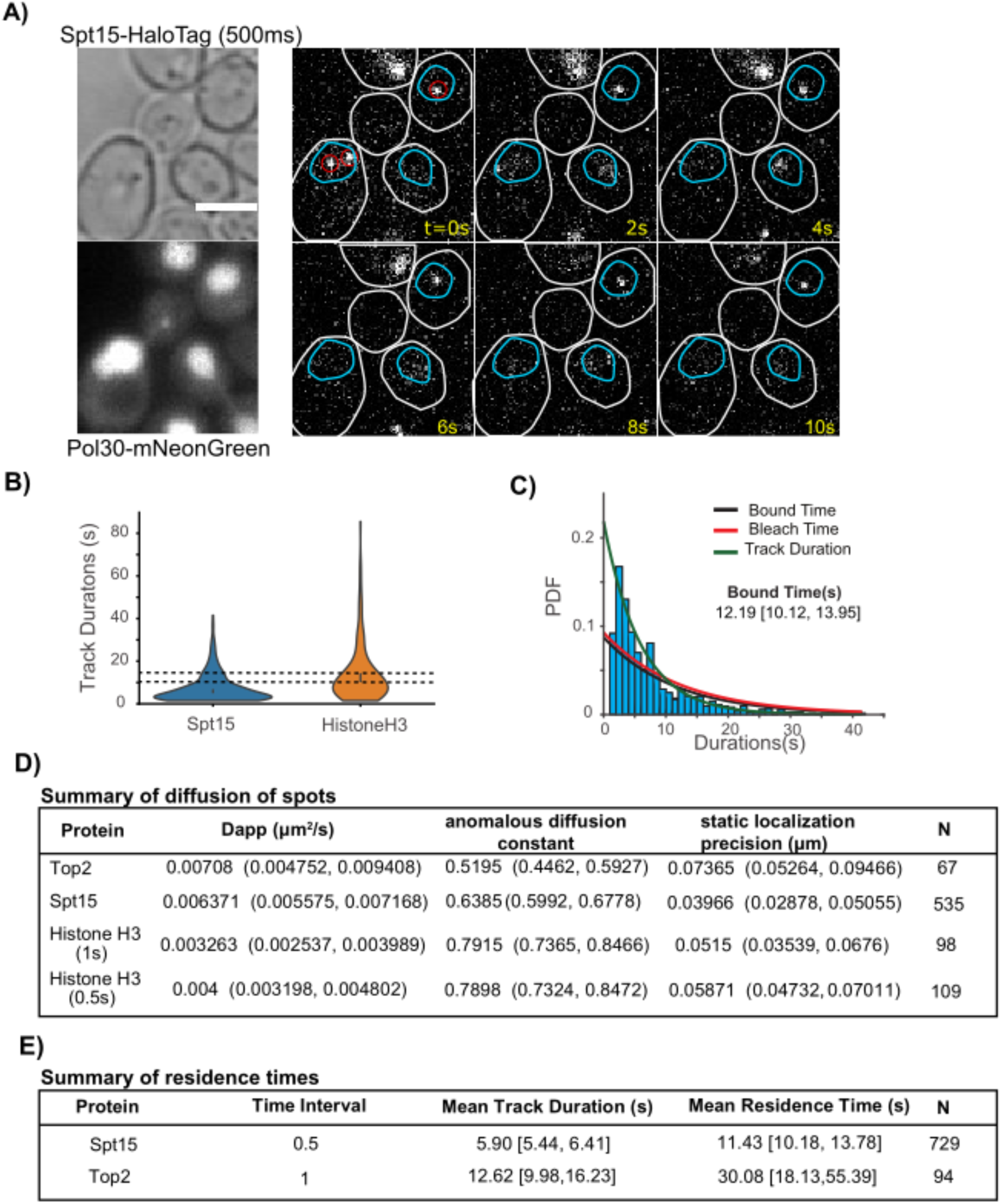
Estimation of residence time for TBP. A) Example of SPT15-HaloTag (TBP) timelapse collected with continuous exposure, after photoactivation. Red circles indicate molecules classified as being bound by Bound2Learn. Also shown is the SME of a Pol30-mNeonGreen z- stack. Scale bar = 3um. B) Violin plots of track durations for TBP (N = 729), and Histone H3 (N = 242). Error bars represent 95% confidence intervals. Dotted horizontal lines represent upper and lower bounds of the 95% confidence interval of Histone H3. C) Histogram and fit on TBP track durations, along with estimate for bound time. D) Estimates for diffusive properties and static localization errors of ML classified bound tracks, obtained through fitting averaged MSD curves. E) Summary of results showing both mean track duration estimates and mean bound time estimates from combined data sets. 95% confidence intervals are shown next to estimates.

To increase our confidence that the tracks classified as being DNA bound were actually DNA-bound, we tested if they had diffusive properties similar to chromosomal loci. We estimated the apparent diffusion coefficients and anomalous diffusion constants for the different budding yeast proteins used here, by fitting to the τ vs MSD (averaged over tracks) curve (**Materials and Methods**) (36). Remarkably, the estimates were not only consistent among the different proteins (Dapp ranging from 0.003263um^2^/s to 0.007080um^2^/s) but were consistent with previously estimated values of chromosomal loci in budding yeast of Dapp ∼0.0025um^2^/s and α ∼ 0.5 (20) (**Figure 5D**). We did notice that histone H3 had slightly different values compared to TBP and Top2, but given that histones are widespread on chromosomes, it is possible that we detected histone H3 molecules bound to chromosomal segments where TBP and Top2 do not bind frequently (e.g. telomeres) (36).

The results from budding yeast, summarized in **Figure 5E,** show estimates for mean track durations and mean bound times for TBP and Top2. Overall, these results show that our approach can accurately determine residence times across a range of binding behaviours in two very different organisms.

## Discussion

Here we have provided a robust, easy-to-use classification approach to isolate tracks of DBP and estimate residence times from long-capture single-molecule data. Commonly used approaches to classify bound molecules by applying thresholds on various track traits (diffusion coefficient, step- size per frame of DBP, PSF, etc.) often are not sufficient for rigorous classification, require additional analysis steps and/or experiments, or may discard too many tracks (1,2,18,24). Our approach bypass these limitations.

The main limitation of our approach is that is still dependent on good image quality, despite performing better than previous approaches. Lower intensity will not only lead to lower quality values but also overestimate speed variables for classification due to increased localization error. Similarly, the classification error in our approach will increase when the diffusive and DNA-bound fractions have similar mobility, although this is likely true for most classifiers. Another limitation is that constructing training data sets manually can take significant time and choosing the right parameter values for building random forest models requires some expertise. However, our results suggest that once the initial models have been created, they can be used across a range of conditions whether that be different exposure times, time intervals, data quality, and are amenable for use in a high-throughput manner. During this study, we had to adjust our simulation parameters (akin to adjusting experimental conditions) to detect two binding regimes in cases where there was a heterogeneous population of bound molecules, similar to what a recent study has reported (28). Nonetheless, this is not an issue with our classification approach but rather the acquisition settings used (or in our case, simulation parameters), and our results suggest simulations can help identifying the optimal acquisition conditions to use for detecting multiple binding states.

Use of Bound2Learn solves some of the limitations with previous approaches for classification of DNA-bound molecules, demonstrating an advantage on using ML approaches. Future development of this approach should focus on decreasing its sensitivity to image quality. However, we expect that Bound2Learn will improve in the analysis of other DNA-binding proteins, and other proteins interacting to relatively immobile structures in the cell. We also expect that this approach will be easily applicable to other experimental models, including mammalian cells.

## Data and materials availability

All data, code, and materials used in this work can be found at https://github.com/Reyes-LamotheLab/Bound2Learn.

## Acknowledgements

We thank Tatiana Karpova (NCI) for providing the yeast strains carrying H3-Halo and pdr5Δ;Luke Lavis (Janelia Farms) for providing the Halo substrate coupled to PA- JF549 dye.

## Author contributions

NK, Conceptualization, Formal analysis, Investigation, and Writing. ZWEH, Formal analysis, Investigation, RR-L, Supervision, Funding acquisition.

