## SupplementaryInformation for "Bound2Learn: A Machine Learning Approach for Classification of DNA-Bound Proteins from Single-Molecule Tracking Experiments"

### Supplementary Information

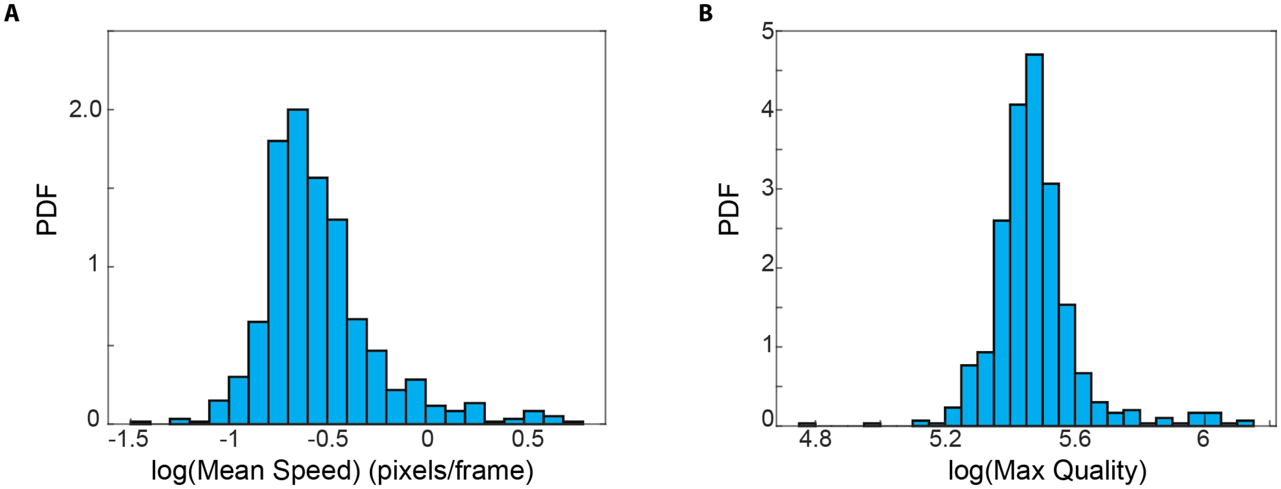

**Figure S1-** A) Representative distribution of the log(mean speed) values for tracks classified as being bound from training data set (500ms, *E.coli*). B) – Representative distribution of the log(maximum quality) values for tracks classified as being bound from training data set (500ms, *E.coli*).

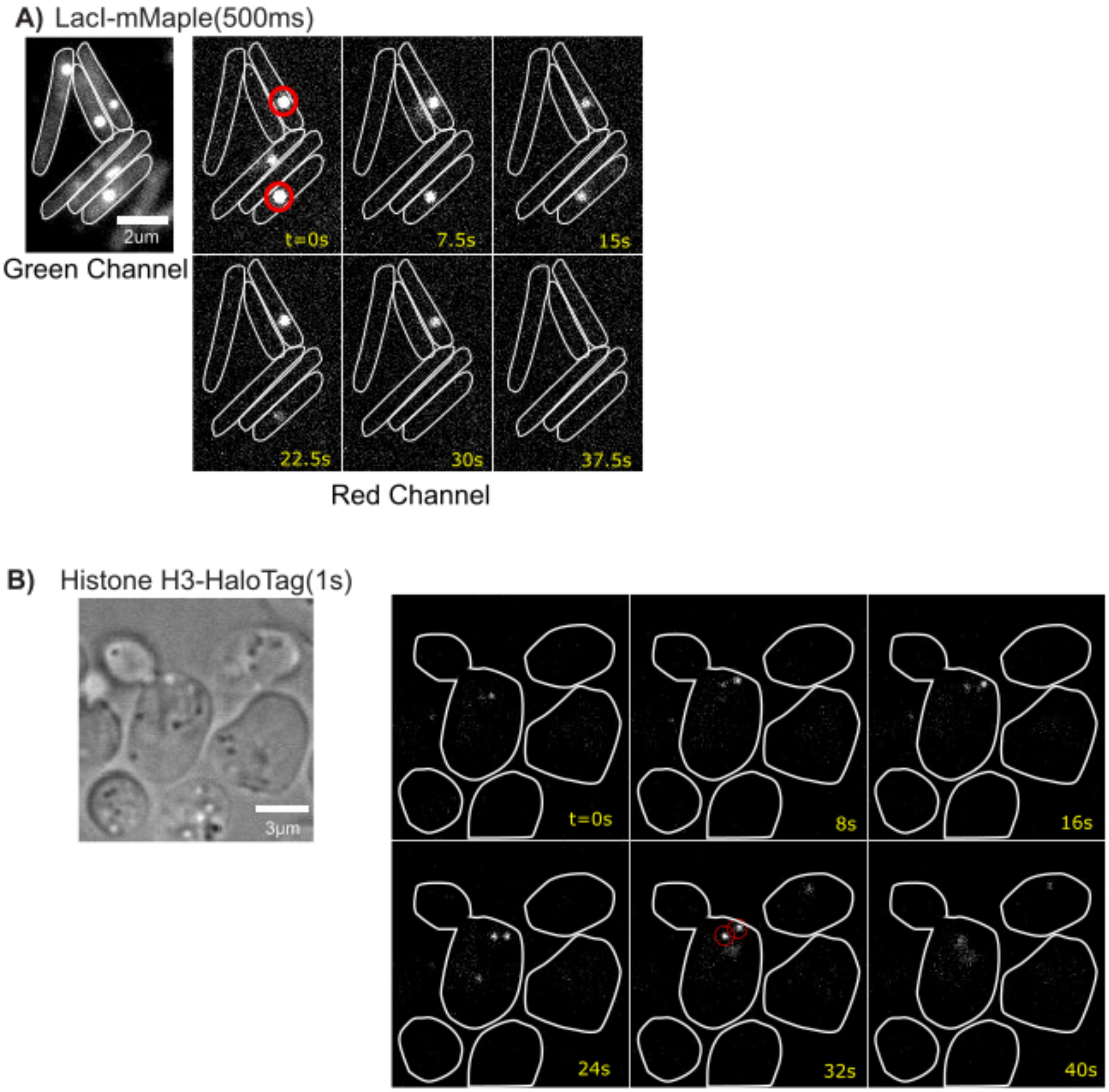

**Figure S2-** A) Example of timelapse for LacI-mMaple collected with continuous exposure acquisition. B) Example of timelapse for Histone H3-HaloTag collected with 500ms exposure, and 1s time interval acquisition. Red circles indicate molecules classified as being bound by Bound2Learn.

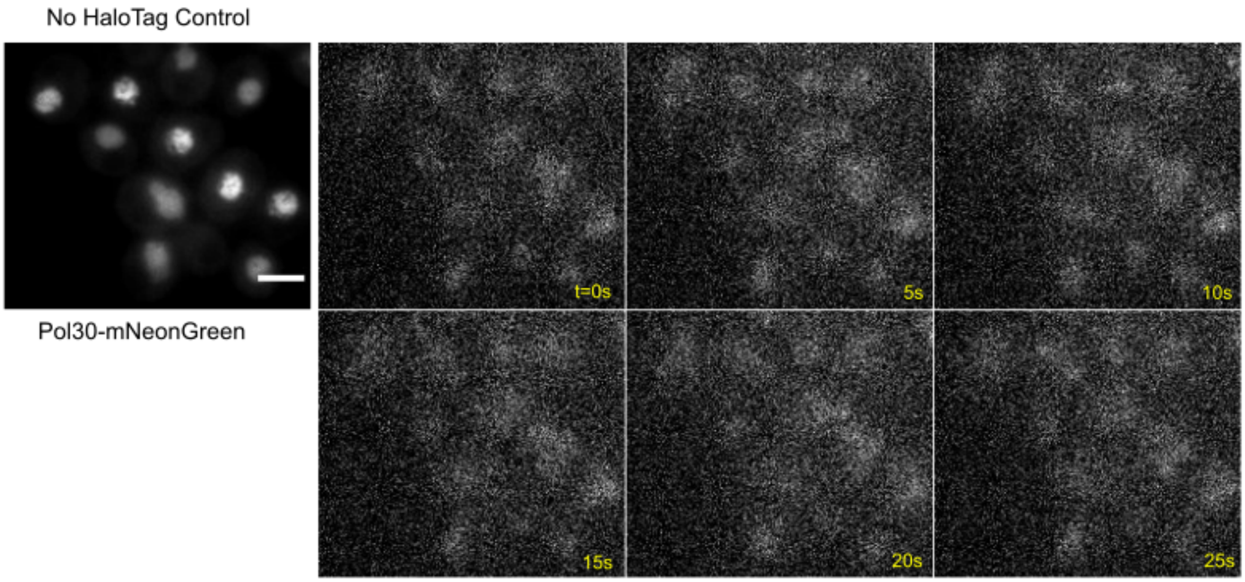

**Figure S3-** Example of timelapse for no HaloTag control (strain ZEY098), which had no HaloTagged protein but still had Pol30-mNeonGreen. Scale bar = 3um. Data collected with continuous exposure acquisition.

| Training Data | Number of Trees | Minimum Leaf Size | Predictors to Sample at Node | Bag Fraction | OOB error | Number of Tracks |
|---|---|---|---|---|---|---|
| ***E.coli* (simulation)** | | | | | | |
| 500ms (ML model 1) | 6000 | 50 | 2 | 0.5 | 0.034 | 1263 |
| 500ms (ML model 2) | 6000 | 50 | 2 | 0.5 | 0.0079 | 1263 |
| 100ms (ML model 1) | 6000 | 50 | 2 | 0.5 | 0.0195 | 1075 |
| 100ms (ML model 2) | 6000 | 50 | 2 | 0.5 | 0.014 | 1075 |
| **Budding Yeast (simulation)** | | | | | | |
| 500ms (ML model 1) | 10000 | 50 | 2 | 0.8 | 0.03 | 967 |
| 500ms (ML model 2) | 10000 | 50 | 2 | 0.8 | 0.0269 | 967 |
| **LacI** | | | | | | |
| 500ms (ML model 1) | 10000 | 70 | 2 | 0.5 | 0.1634 | 1438 |
| 500ms (ML model 2) | 10000 | 70 | 2 | 0.5 | 0.0709 | 1438 |
| **Histone H3** | | | | | | |
| 500ms, 1s time interval (ML model 1) | 6000 | 50 | 2 | 0.5 | 0.1151 | 1251 |
| 500ms, 1s time interval (ML model 2) | 6000 | 50 | 2 | 0.5 | 0.036 | 1251 |

**Table S1.** Parameter values used to construct random forests.

| Training Data | Tbleach (s) | Tbound(s) | Tsearch(s) | $D_{mobile}$ (um^2/s) | $D_{bound}$ (um^2/s) | Mobile fraction | Bound Fraction | Integrated Spot Intensity |
|---|---|---|---|---|---|---|---|---|
| **Ecoli** | | | | | | | | |
| 500ms exposure (no time interval) | 10 | 100 | 100 | 0.5 | 0.005 | 0.3 | 0.7 | 3000 |
| 100ms exposure (no time interval) | 2 | 100 | 100 | 0.5 | 0.005 | 0.3 | 0.7 | 3000 |
| **Budding Yeast** | | | | | | | | |
| 500ms exposure (no time interval) | 10 | 100 | 100 | 0.5 | 0.005 | 0.3 | 0.7 | 3000 |
| **Experimental** | | | | | | | | |
| **Ecoli** | | | | | | | | |
| 500ms exposure (1s time interval) | 20 | 8 | 10000000 | 0.5 | 0.005 | 0.5 | 0.5 | 3000 |
| 500ms exposure (1s time interval, lower spot intensity) | 20 | 8 | 10000000 | 0.5 | 0.005 | 0.5 | 0.5 | 2000 |
| 100ms exposure (no time interval) | 2 | 1 | 10000000 | 0.5 | 0.005 | 0.7 | 0.3 | 3000 |
| 100ms exposure (no time interval, higher $D_{mobile}$ ) | 2 | 1 | 10000000 | 5 | 0.005 | 0.7 | 0.3 | 3000 |
| 100ms exposure (no time interval, mixed bound population) | 10 | 1s/7s | 10000000 | 0.5 | 0.005 | 0.1 | 0.45/0.45 | 3000 |
| **Budding Yeast** | | | | | | | | |
| 500ms exposure (1s time interval) | 20 | 8 | 10000000 | 0.5 | 0.005 | 0.5 | 0.5 | 3000 |
| 500ms exposure (1s time interval, lower spot intensity) | 20 | 8 | 10000000 | 0.5 | 0.005 | 0.5 | 0.5 | 2000 |

**Table S2 –** Parameters used for simulations of training data and experimental data.

| **E.coli** | **Spot Intensity = 3000** | **Spot Intensity = 2000** |
| --- | --- | --- |
| **1s Interval (500ms exposure)** | | |
| Bound Time [95% Confidence Interval] | 6.76[5.41, 8.59] | 7.21[5.65, 8.95] |
| Number of tracks predicted to be bound | 169 | 159 |
| Prediction Accuracy | 0.93 | 0.99 |
| Recovery Error | 0.11 | 0.11 |
| **100ms Exposure (no time interval)** | | |
| Bound Time [95% Confidence Interval] | 0.97[0.75, 1.21] | |
| Number of tracks predicted to be bound | 156 | |
| Prediction Accuracy | 0.85 | |
| Recovery Error | 0.18 | |
| **100ms Exposure (no time interval). Predicted using 500ms Training Data** | | |
| Bound Time [95% Confidence Interval] | 0.96[0.75, 1.22] | |
| Number of tracks predicted to be bound | 158 | |
| Prediction Accuracy | 0.86 | |
| Recovery Error | 0.16 | |
| **100ms Exposure (no time interval). $D_{mobile}$ = 5um^2/s** | | |
| Bound Time [95% Confidence Interval] | 0.90[0.71, 1.13] | |
| Number of tracks predicted to be bound | 142 | |
| Prediction Accuracy | 0.98 | |
| Recovery Error | 0.08 | |
| **Diffusion Coefficient Analysis - 1s Interval (500ms exposure)** | | |
| Bound Time [95% Confidence Interval] | 3.72[3.05,4.46] | 5.06 [3.97, 6.33] |
| Number of tracks predicted to be bound | 291 | 203 |
| Prediction Accuracy | 0.54 | 0.79 |
| Recovery Error | 0.12 | 0.096 |
| **Budding Yeast** | | |
| Bound Time [95% Confidence Interval] | 7.26[5.97, 8.77] | 7.17[5.92, 8.82] |
| Number of tracks predicted to be bound | 232 | 227 |
| Prediction Accuracy | 0.97 | 0.97 |
| Recovery Error | 0 | 0 |

**Table S3.** Results for simulated experimental data.

| **Protein** | **Time Interval** | **Mean Track Duration (s)** | **N** |
|---|---|---|---|
| LacI | 0.5 | 15.09 (13.42, 16.75) | 316 |
| LacI | 0.5 | 10.91 (8.74, 13.64) | 129 |
| ε | 1 | 7.26 (6.08, 8.84) | 135 |
| ε | 1 | 7.74 (6.14, 9.96) | 78 |
| β | 1 | 18.55 (16.14, 20.95) | 229 |
| β | 1 | 15.19 (12.87, 19.52) | 160 |
| β | 5 | 45.47 (40.36, 52.28) | 279 |
| β | 5 | 48.51 (41.07, 56.72) | 158 |
| Histone H3 | 0.5 | 12.19 [10.71,13.87] | 242 |
| Histone H3 | 1 | 21.74 [18.41, 26.50] | 123 |
| TBP | 0.5 | 6.18 [5.55,6.79] | 495 |
| TBP | 0.5 | 5.32[4.61, 6.07] | 234 |
| Top2 | 1 | 11.49 [7.77, 19.05] | 43 |
| Top2 | 1 | 13.57[10.69, 17.62] | 51 |

**Table S4.** Results with *E.coli* and budding yeast data. The 95% confidence are presented next to the estimates.

| Strain | Genotype |
|---|---|
| **BY4741** | MATa his3Δ1 leu2Δ0 met15Δ0 ura3Δ0 |
| **BY4742** | MATα his3Δ1 leu2Δ0 lys2Δ0 ura3Δ0 |
| **BY4743** | MATa/α his3Δ1/his3Δ1 leu2Δ0/leu2Δ0 LYS2/lys2Δ0 met15Δ0/MET15 ura3Δ0/ura3Δ0 |
| **YTB31** | MATα his3Δ1 leu2Δ0 lys2Δ0 ura3Δ0 POL30-mNeonGreen-Nat |
| **YTK1414** | MATa/α his3Δ1/his3Δ1 leu2Δ0/leu2Δ0 LYS2/lys2Δ0 met15Δ0/MET15 ura3Δ0/ura3Δ0 PDR5/pdr5Δ::KanMX |
| **ZEY098** | MATa his3Δ1 leu2Δ0 LYS2 met15Δ0 ura3Δ0 pdr5Δ0::KanMX POL30-mNeonGreen-Nat |
| **ZEY075** | MATa his3Δ1 leu2Δ0 LYS2 met15Δ0 ura3Δ0 pdr5Δ0::KanMX POL30-mNeonGreen-Nat TOP2-Halo-HygB |
| **ZEY157** | MATα his3Δ1 leu2Δ0 lys2Δ0 MET15 ura3Δ0 pdr5Δ0::KanMX POL30-mNeonGreen-Nat SPT15-Halo-HygB |
| **YTK1434** | MATa his3Δ1 leu2Δ0 met15Δ0 ura3Δ0 pdr5Δ0::KanMX HHT1-Halo-URA3 |

**Table S5.** Strains used**.** The *S. cerevisiae* strains used in this study are listed along with their genotypes.

| Primer | Description | Sequence |
|---|---|---|
| **TB81** | C-terminal mNeonGreen tagging of PCNA (F) | cctacagtttttcttggctcctaaatttaatgacgaagaaGGTGACGGTGCTGGTTTA |
| **TB82** | C-terminal mNeonGreen tagging of PCNA (R) | tttattatttttagtatacaactatatagataatttacatCACAGGAAACAGCTATGACC |
| **TB98** | Screen C-terminal tag of PCNA (F) | AGAGTTGGTATCAGGCTCTC |
| **TB99** | Screen C-terminal tag of PCNA (R) | AAGCTGATATTTAACGCATCTTAG |
| **TOP2insF** | C-terminal Halo tagging of Top2 (F) | aggaaaaccaaggatcagatgtttcgttcaatgaagaggatGGTGACGGTGCTGGTTTA |
| **TOP2insR** | C-terminal Halo tagging of Top2 (R) | acatataaaaagaatggcgctttctctggataaatattatCACAGGAAACAGCTATGACC |
| **TOP2seqF** | Screen C-terminal tag of Top2 (F) | ACTATCTGGTGAAAGCGACC |
| **TOP2seqR** | Screen C-terminal tag of Top2 (R) | ACGATGTTTTTCGCCCAGGC |
| **NK46** | C-terminal Halo tagging of Spt15 (F) | tgaagctatataccctgtgctaagtgaatttagaaaaatgGGTGACGGTGCTGGTTTAAT |
| **NK47** | C-terminal Halo tagging of Spt15 (R) | aatagaaaacctttttttcttttcgtctactccttccccaCAGTATAGCGACCAGCATTC |
| **NK50** | Screen C-terminal tag of Spt15 (F) | CTCCTATGAGCCAGAATTG |
| **NK51** | Screen C-terminal tag of Spt15 (R) | CTCCTATGAGCCAGAATTG |

**Table S6.** Primers used in this study, with a short description and their sequence.
